# Extracellular electron transfer may be an overlooked contribution to pelagic respiration in humic-rich freshwater lakes

**DOI:** 10.1101/392027

**Authors:** Shaomei He, Maximilian P. Lau, Alexandra M. Linz, Eric E. Roden, Katherine D. McMahon

## Abstract

Humic lakes and ponds receive large amounts of terrestrial carbon and are important components of the global carbon cycle, yet how their redox cycling influences the carbon budget is not fully understood. Here we compared metagenomes obtained from a humic bog and a clearwater eutrophic lake, and found a much larger number of genes that might be involved in extracellular electron transfer (EET) for iron redox reactions and humic substance (HS) reduction in the bog than in the clearwater lake, consistent with the much higher iron and HS levels in the bog. These genes were particularly rich in the bog’s anoxic hypolimnion, and were found in diverse bacterial lineages, some of which are relatives of known iron oxidizers or iron/HS reducers. We hypothesize that HS may be a previously overlooked electron acceptor and EET-enabled redox cycling may be important in pelagic respiration and greenhouse gas budget in humic-rich freshwater lakes.

## INTRODUCTION

Inland lakes receive allochthonous carbon (C) fixed in their catchment areas, and play an important role in the cycling of terrestrial C and affect global C budgets. Many northern freshwater lakes are experiencing a “browning” process, and this trend may continue with changes in precipitation patterns and atmospheric deposition chemistry (1–3). A leading factor contributing to the brownification is the increasing inputs of allochthonous dissolved organic C (DOC) (4). A major component of terrestrially derived allochthonous DOC in freshwater is humic substances (HS), which are heterogeneous mixtures of naturally occurring recalcitrant organic carbon derived from plant and animal decay (5). Another factor contributing to surface water brownification is increasing iron (Fe) inputs (6), which were positively correlated to the increasing organic C inputs (7, 8). This correlation may be partly due to the complexation of Fe by organic matter, in particular HS, as the complexation may increase Fe leaching from catchment soil and maintain Fe in the water column instead of sedimentation within the receiving water body (8, 9).

Understanding the roles of HS and Fe in freshwater lakes is critical to predict how such ecosystems will respond to the browning process, and to more accurately dissect overall lake “metabolism” (10, 11). Humic lakes feature intensively brown-colored waters originating from high concentration in HS and Fe. As a redox-active element, Fe plays a role in defining redox conditions and C cycling; yet the complex roles of HS have not been fully recognized, despite its high concentrations in humic lakes. On one hand, HS and the more labile low-molecular weight C derived from the photodegradation of HS is an important C source and electron donor for heterotrophic respiration in humic lakes (12, 13). On the other hand, HS has also been recognized as an electron acceptor through the reduction of their quinone moieties (14, 15), and its electron accepting capacity (EAC) is fully regenerable under recurrent anoxic conditions (16). However, most prior research on the EAC of HS considered the impact on C-cycling in wetlands, sediments and soils, rather than truly pelagic ecosystems (16–19). Recently, a study on a humic lake showed that native organic matter with more oxidized quinone moieties and therefore higher EAC favored freshwater bacterial growth and production under anoxic conditions, and further suggested organic matter as an important electron acceptor in stratified lakes with oxycline fluctuations (20). Despite this, the role of HS as an electron acceptor in freshwater lakes has not been widely appreciated.

Theoretically, if HS is used to respire organic C, it has the potential to lower methane emissions from lakes. The reduction potential distribution in HS suggests HS reduction to be thermodynamically more favorable than methanogenesis in anoxic waters (16). As the resulting competitive mitigation of methanogenesis was observed in peat bogs and peat soils (21–23), a similar process is expected for pelagic respiration in lakes. Therefore, we judge it timely to further explore the contribution of HS and Fe reduction to pelagic respiration in freshwater lakes.

In humic lakes, light does not penetrate deep into the water column due to the absorbance by HS and Fe. Therefore, humic lakes generally have a shallower phototrophic (and therefore oxygenated) zone than clearwater lakes during stratification (13), leaving a larger proportion of water column under anoxic conditions. Due to this redox distribution and their high concentrations, HS and Fe may become important terminal electron acceptors in humic lakes. Thus, here we present the hypothesis that HS and Fe redox cycling is more significant in humic lakes than in clearwater lakes, and that these redox processes may influence ecosystem-level C budgets (i.e. overall lake metabolism). As a preliminary examination of this hypothesis, we studied two contrasting temperate lakes, including a small humic lake, Trout Bog, in which the DOC is primarily of terrestrial origin; and a large eutrophic clearwater lake, Lake Mendota, which has much lower concentrations of HS and Fe than Trout Bog, with most of its DOC being produced in-lake via photosynthesis. Detailed lake characteristics are listed in **Table S1**. Three combined assemblies of time-series metagenome libraries previously obtained from Lake Mendota epilimnion (ME), Trout Bog epilimnion (TE), and Trout Bog hypolimnion (TH), respectively, and over 200 metagenome-assembled genomes (MAGs) were recovered from these combined assemblies (21). We examined these metagenomes and MAGs to identify genes involved in HS and Fe redox processes to compare their distributions in the two contrasting lakes.

Due to the high molecular weight of HS and the poor solubility of Fe(III), these electron acceptors are reduced extracellularly via a process called extracellular electron transfer (EET). The reduced HS and Fe can be abiotically re-oxidized by oxygen under oxic conditions. In addition, biological Fe(II) oxidation may also occur, which employs EET due to the poor solubility of the reaction product, Fe(III). One form of oxidoreductase in Fe redox EET processes involves outer-surface proteins, such as Cyc2, a monoheme cytochrome *c* typically found in Fe(II) oxidizers (22–24), and multiheme *c*-type cytochromes (MHCs) in Fe(III) reducers (25, 26). Another form of EET oxidoreductase forms a “porin-cytochrome *c* protein complex” (PCC) (27), in which the oxidoreductase, usually a MHC, is secreted to the periplasm and embedded into a porin on the outer membrane to form the EET conduit. Most Fe(III) reducers can also reduce HS (28, 29), and probably use the same EET systems to transfer electrons to HS. For example, in *Geobacter sulfurreducens*, a number of outer membrane MHCs that are important in the reduction of Fe(III) are able to reduce extracellular AQDS and HS (30), and in *Shewanella oneidensis*, the porin and periplasmic MHC components of its Fe(III)-reducing PCC are essential for AQDS and HS reduction (31, 32). These findings suggest that reduction of the quinone moieties in HS is a non-specific redox process by EET systems.

In this study, we searched for putative EET genes (including PCC, outer surface MHCs not associated with PCC, and Cyc2) in MAGs and metagenomes to examine if these genes are indeed more abundant in the humic bog than in the clearwater lake. Method details on the identification and quantification of putative EET genes were described in Supplemental Methods. All (meta)genome data are publicly available at the Integrated Microbial Genomes & Microbiomes (IMG/M, https://img.jgi.doe.gov/m). The IMG IDs for the ME, TE, and TH metagenomes are 3300002835, 3300000439, and 3300000553, respectively, and the IMG IDs for putative EET gene-containing MAGs (together with details on these MAGs and putative EET genes in the three metagenomes) are listed in **Tables S2** and **S3**.

## RESULTS

MHCs are important components of EET systems involved in Fe redox reactions and HS reduction. In particular, MHC with large numbers of hemes may be able to form molecular “wires” for conducting electrons from the periplasmic space across the outer membrane (33, 34). We therefore estimated the normalized abundance of MHCs with at least five heme-binding sites in the metagenomes. In general, TH had the highest abundance of MHCs, followed by TE and ME, and such differences were even more pronounced for MHCs with at least eight heme-binding sites (**Fig. 1A**). Some of these MHCs are components of other redox enzyme complexes, such as the pentaheme and hexaheme MHCs in alternative complex III (ACIII), and octaheme MHCs in tetrathionate reductases and hydroxylamine oxidoreductases. Putative EET MHC components (i.e. MHCs in PCC and outer surface MHCs not associated with PCC, as listed in **Table S2**) were much more frequently found in MHCs with large heme binding sites (e.g. >9), and these putative EET genes were more abundant in TH than TE, and nearly absent in the ME metagenome (**Fig. 1B**). This may indicate that MHC-based EET potential was more significant in the anoxic layer than in the oxic layer of the humic bog, and was minimal in the oxic layer of the clearwater lake with low Fe and HS concentrations. Notably, the largest number of heme-binding sites (i.e. 51) was found in an MHC component of a putative PCC, encoded in an un-binned contig in the TE metagenome (**Table S2**).

**Fig. 1.**
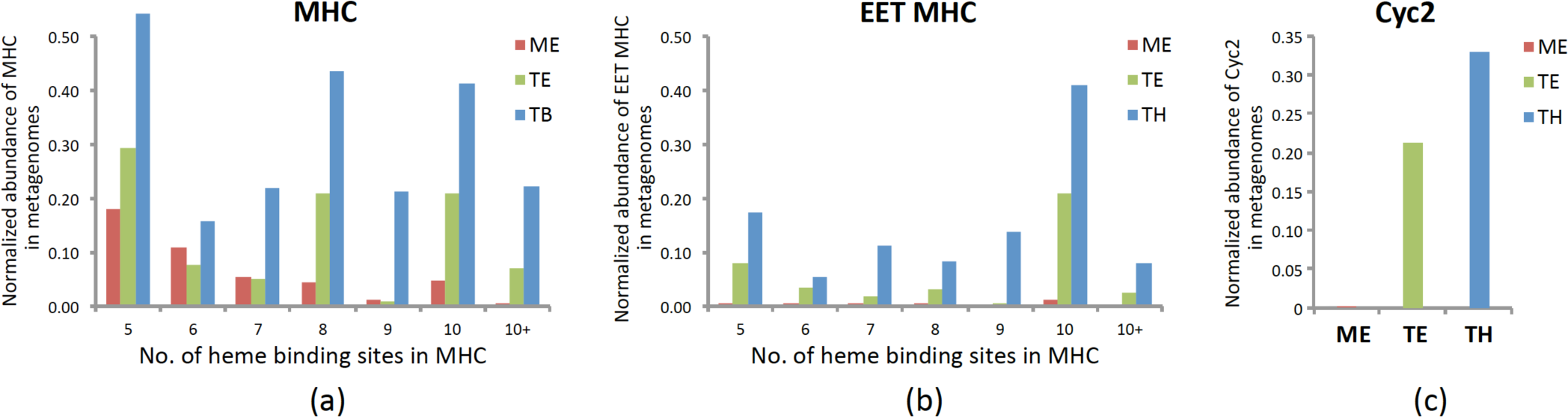
Normalized abundance of multiheme *c*-type cytochromes (MHCs) (a), MHCs with putative EET functions (i.e. MHCs in PCC and outer surface MHCs not associated with PCC) (b), and Cytochrome Cyc2 homologs (c) found in metagenomes obtained from Lake Mendota’s epilimnion (ME), and Trout Bog’s epilimnion (TE) and hypolimnion (TH), respectively. In (a) and (b), normalized abundance was reported for MHCs with 5 to 10, and > 10 heme binding sites respectively. The normalized abundance was obtained by mapping metagenome reads to assembled contigs and the read coverage was then normalized by the average read coverage of single-copy conserved bacterial housekeeping genes in the same metagenome. See Supplemental Methods for details on the calculation of normalized abundance.

### Porin-cytochrome *c* protein complex (PCC) genes

The best studied PCC system, MtrABC (consisting of a porin, a periplasmic decaheme Cyt *c*, and an extracellular decaheme Cyt *c*), was first identified in *S. oneidensis* as being essential for Fe(III) reduction (35). Their homologous PCCs, PioAB and MtoAB, which lack the extracellular MHC component, were suggested to be involved in Fe(II) oxidation in the phototrophic *Rhodopseudomonas palustris* TIE-1 (36) and the microaerophilic Fe(II) oxidizers in the family of *Gallionellaceae* (37–39), respectively. The more recently discovered PCC proteins in *G. sulfurreducens* are not homologous to MtrABC, but are also encoded in operons with genes encoding a porin (OmbB), a periplasmic octaheme Cyt *c* (OmaB), and an outer-membrane dodecaheme Cyt *c* (OmcB) (40). This suggests that multiple PCC systems evolved independently, and may provide a clue to search for new types of PCC by examining genome-level organization. For example, putative novel PCC genes not homologous to previously identified PCCs were found in some Fe(II) oxidizer genomes by searching for the unique genetic organization of porin-and periplasmic MHC-coding genes (41).

Nearly all MtrAB/MtoAB/PioAB homologs were recovered in Trout Bog, and mostly from TH (Table S2). They are present in MAGs affiliated with the Proteobacteria, including the Fe(II)-oxidizing *Gallionella* and *Ferrovum*, Fe(III)-reducing *Albidiferax*, Fe(III)- and AQDS-reducing *Desulfobulbus* and genera not known for EET, such as *Polynucleobacter*, *Desulfocapsa*, and *Methylobacter* (**Fig. 2**). Interestingly, among the 46 *Polynucleobacter* genomes available at IMG/M (https://img.jgi.doe.gov/m), MtrAB/MtoAB/PioAB homologs were only found in *Polynucleobacter* recovered from a wetland and two humic lakes (including Trout Bog and Lake Grosse Fuchskuhle located in Brandenburg, Germany), suggesting that this PCC might be an acquired trait of some *Polynucleobacter* spp. adapting to humic-rich environments.

**Fig. 2.**
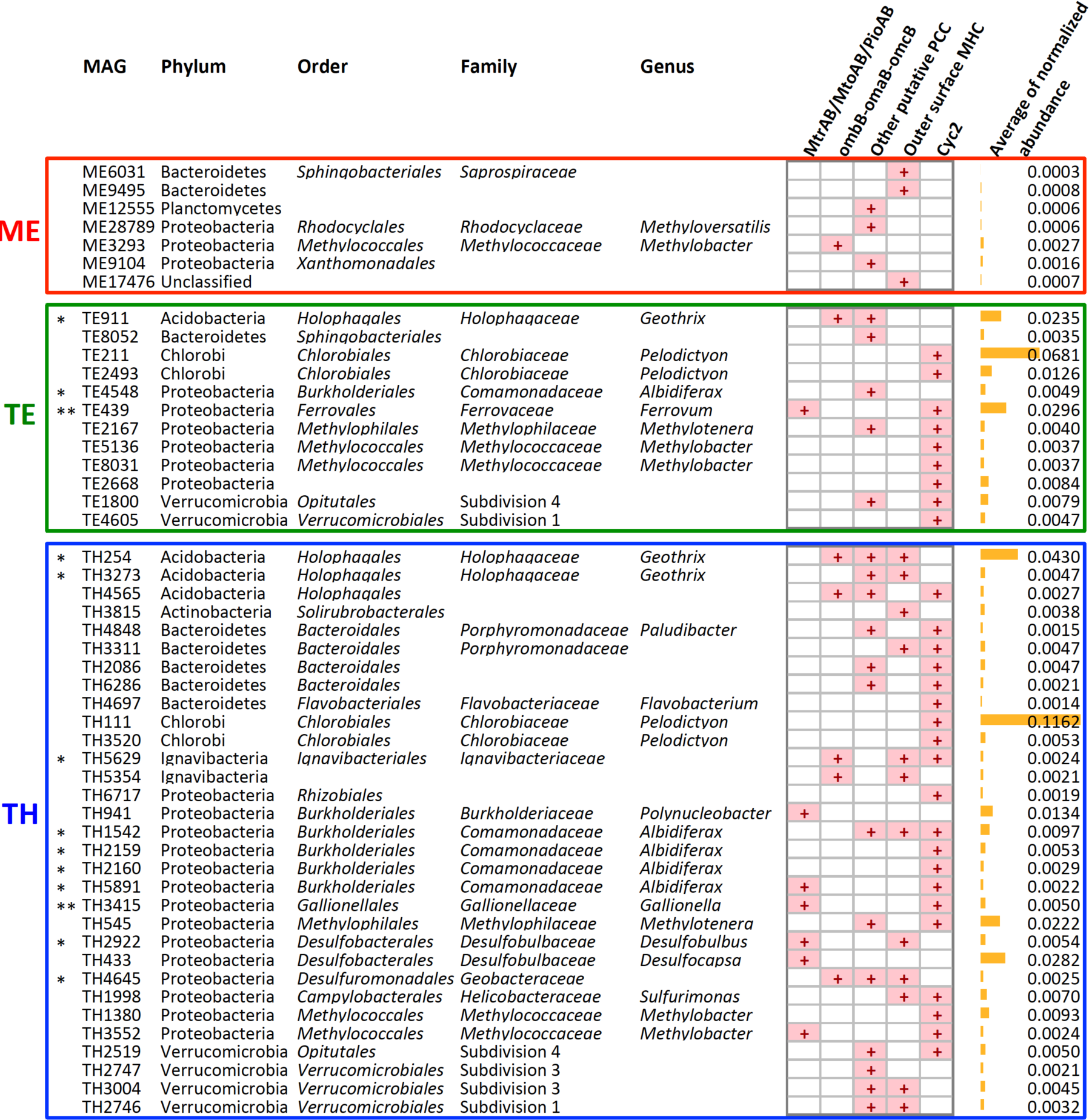
Occurrence of putative EET genes in MAGs and their normalized abundance in the metagenome as measured by mapping reads to assembled contigs for read coverage and normalizing by the average coverage of single-copy conserved bacterial housekeeping genes (see Supplemental Methods for details). * indicates MAGs with Fe(III) reducing relatives, and ** indicates MAGs with Fe(II) oxidizing relatives. The presence of putative EET genes was indicated with “+”.

Homologs of another studied PCC (represented by OmbB-OmaB-OmcB in *Geobacter* spp.) were present in MAGs affiliated with relatives of known Fe(III) (and HS) reducers, including *Geothrix*, *Ignavibacteriaceae* and *Geobacteraceae*, as well as in *Methylobacter* (**Fig. 2**).

Based on the unique genetic organization of PCC-encoding genes, we found a number of putative PCC that do not share a significant sequence homology with known PCCs, probably representing novel PCC types. These putative PCC genes were present in Fe(III) (and HS) reducers (*Geothrix*, *Albidiferax*, and *Geobacteraceae*) and bacteria not known for EET, including *Methylotenera*, *Methylobacter*, *Methyloversatilis*, and a number of Bacteroidetes and Verrucomicrobia (**Fig. 2**). Among them, Verrucomicrobia with putative PCC genes were previously found in humic-rich environments, such as soils and lake sediment, in addition to the Verrucomicrobia MAGs from Trout Bog (42).

### Outer surface MHCs not associated with PCC

Outer surface MHCs that are not PCC components may also be involved in EET. Examples include the OmcE, OmcS, and OmcZ in *G. sulfurreducens* (25, 26), outer surface MHCs in Gram-positive Fe(III)- and AQDS-reducing Firmicutes (43), and MHCs in deltaproteobacterial sulfate-reducing bacteria that may be responsible for EET with its anaerobic CH_4_-oxidizing archaeal syntrophic partner (44).

Here, we found a number of non-PCC-associated outer surface MHCs in the metagenomes (**Table S2**) and MAGs, including Fe(III)- (and HS-) reducing taxa (*Albidiferax*, *Geothrix*, *Desulfobulbus*, *Ignavibacteriaceae* and *Geobacteraceae*), and several members in the Bacteroidetes and Verrucomicrobia phyla (**Fig. 2**). In particular, seven genes predicted to encode MHCs located on the cell wall were found in a Gram-positive actinobacterial MAG classified to *Solirubrobacterales* from TH, and four of these genes are located in the same gene cluster with up to 15 heme-binding sites in a single MHC (**Table S2**), probably involved in electron transfer on the cell wall.

### Cytochrome Cyc2

Cyc2 is an outer membrane *c*-type cytochrome with one heme-binding motif in the N-terminus and a predicted porin structure at the C-terminus, and was therefore proposed as a fused PCC (45). Cyc2 was originally identified as the Fe(II) oxidase in acidiphilic *Acidithiobacillus ferrooxidans* (23) with distant homologs later found in neutrophilic microaerobic *Mariprofundus* spp. (22) and some other neutrophilic Fe(II) oxidizers (24, 41).

Similar to EET MHC genes, the normalized abundance of total Cyc2-like genes was much higher in the TH than in the TE metagenome, and Cyc2-like genes were largely absent in the ME metagenome (**Fig. 1C**). Cyc2 homologs were present in 29 MAGs exclusively from Trout Bog (**Table S3**), including relatives of Fe(II)-oxidizing genera (*Ferrovum* and *Gallionella*) and Fe(III)- reducing taxa (*Ignavibacteriaceae* and *Albidiferax*), as well as bacteria not known for EET, including *Methylotenera*, *Methylobacter*, *Pelodictyon*, and members in Bacteroidetes and Verrucomicrobia (**Fig. 2**).

## DISCUSSION

With the ongoing brownification of surface water due to increasing inputs of terrestrial C and Fe on a large scale, elucidating the roles and contribution of HS and Fe in redox and C cycling becomes even more relevant to C budgets at an ecosystem level. Here we inspected EET genes/organisms involved in HS and Fe redox processes in two freshwater lakes with contrasting HS and Fe levels to examine if these genes/organisms were more abundant in the humic lake, particularly in its anoxic layer. All together, a total of 103, 36, and 66 MAGs were recovered from the ME, TE, and TH metagenomes, respectively. Among them, putative EET genes were found in 7, 12 and 31 MAGs from ME, TE and TH, respectively (**Fig. 2**). Therefore, a larger fraction of MAGs might encode EET function in Trout Bog, especially in its hypolimnion, than in Mendota. This, together with the normalized abundance of putative EET genes in the three metagenomes (**Fig. 1**), suggests that the genetic potential of EET was more significant in the anoxic layer than in the oxic layer of the humic bog, and was the lowest in the oxic layer of the clearwater lake. This distribution pattern is consistent with the availability of the thermodynamically more favorable electron acceptor, i.e. oxygen, between the two layers and the much higher concentrations of HS and Fe in the bog than in the clearwater lake.

It was not surprising to find putative EET genes in relatives of bacteria that are known to be capable of Fe redox reactions and HS reduction in anoxic lake waters. However, finding putative EET genes in taxa not known for EET functions is intriguing. Like many known EET organisms, some of these bacteria (e.g. Bacteroidetes and Verrucomicrobia) contain multiple sets of putative EET genes. In particular, some *Methylotenera* and *Methylobacter* contain both Cyc2 and putative PCC genes. If these methylotrophs are indeed capable of EET, this may enable insoluble or high-molecular weight substrates, such as Fe(III) and HS, to be used as an electron acceptor to oxidize the methyl-group in methanol and methylamine. Such EET processes, if they occur, could allow methylotrophs to survive in the anoxic layer, and this agrees with the recovery of *Methylotenera* and *Methylobacter* MAGs in the largely anoxic hypolimnion of Trout Bog.

HS have until now usually been more regarded as an electron donor and C source in freshwater lakes, and not as an electron acceptor. However, evidence for the role as an electron acceptor was recently documented in another peat bog lake (20). In the current study, we measured the electron accepting capacity (EAC) of HS in the epilimnion and hypolimnion water of Trout Bog according to Kappler et al. (17), and their EAC was 0.115 and 0.128 mM, respectively (See Supplemental Methods for the determination of lake water EAC). Notably, these values are an order of magnitude higher than the EAC of Fe (~0.01 mM) in Trout Bog. Therefore, HS may be a significant, but previously overlooked source of electron acceptors in this bog system.

Due to its high EAC and concentration, HS may play an important role in the redox cycling in Trout Bog. On one hand, HS facilitates Fe redox reactions by shuttling electrons from Fe(III)-reducers to Fe(III) in heterotrophic respiration (46). On the other hand, HS may be directly used as an electron acceptor to respire the more labile organic C (**Fig. 3**). The anaerobic respiration of organic C with Fe(III) and HS are both thermodynamically more favorable than methanogenesis, therefore promoting the transformation of organic C towards CO_2_, not CH_4_. This may lower the overall global warming potential of greenhouse gas emissions from humic lakes, as CH_4_ is a much more potent greenhouse gas than CO_2_. Because of lake seasonal mixing and more frequent micro-mixing, such as wind-driven turbulence and convectively derived diurnal oxycline fluctuations (20, 47), reduced HS and Fe can be re-oxidized through mixing-introduced oxygenation to regenerate their EAC, which makes these anaerobic respiration processes sustainable in the anoxic layer (**Fig. 3**). In these redox processes, oxygen is the ultimate electron acceptor, and Fe and HS “recharge” the EAC with oxygen for subsequent use when oxygen becomes unavailable in stratified hypolimnia. Hypothetically, such recharging process would increase the effective EAC of humic water and shunt more organic C to anaerobic respiration. Therefore, we hypothesize that HS may be a previously overlooked electron acceptor and EET may be an important contribution to pelagic respiration in humic-rich freshwater lakes. Coupled with C metabolism, EET-enabled HS and Fe redox dynamics can significantly influence C cycling and greenhouse gas emission in humic lakes that experience recurrent oxic-anoxic conditions. The overrepresentation of EET genes/organisms potentially involved in HS and Fe redox processes in the humic lake strongly support this hypothesis, given that the energetic advantage such organisms can obtain stays marginal when powerful recharge mechanisms at the oxic/anoxic interface are lacking. Yet further combined biogeochemical, hydrodynamic, genomic and transcriptomic studies are required to test our hypothesis and reveal organisms and genes actually involved *in-situ*.

**Fig. 3.**
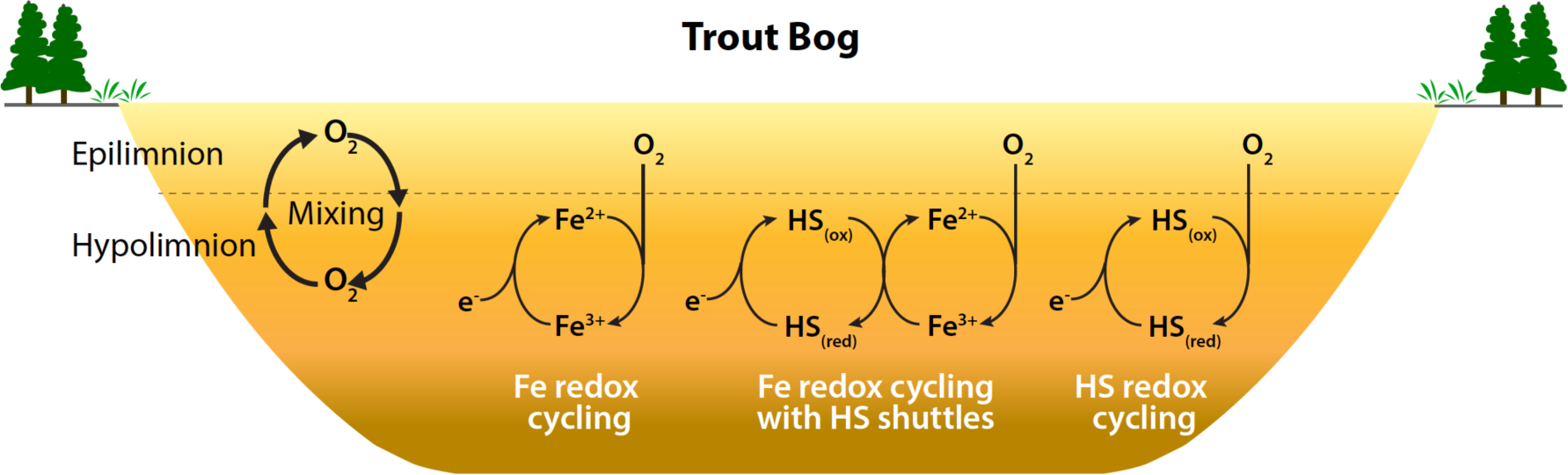
Proposed roles of EET genes in facilitating redox cycling of Fe and HS in the bog. Oxygenation in the hypolimnion through seasonal mixing and more frequent micro-mixing (such as wind-driven turbulence and convectively derived diurnal oxycline fluctuations) regenerates the electron accepting capacity of reduced HS and Fe to enable these anaerobic respiration processes sustainable in the hypolimnion.

## ACKNOWLEDGEMENTS

We thank the North Temperate Lakes Microbial Observatory 2007-2012 field crews, UW-Trout Lake Station, the UW Center for Limnology, and the Global Lakes Ecological Observatory Network for field and logistical support. We give special thanks to past McMahon lab graduate students Ashley Shade, Stuart Jones, Ryan Newton, Emily Read, and Lucas Beversdorf. We acknowledge efforts by many McMahon Lab undergrads and technicians related to sample collection and DNA extraction, particularly Georgia Wolfe. We personally thank the individual program directors and leadership at the National Science Foundation for their commitment to continued support of long term ecological research.

The project was supported by funding from the United States National Science Foundation Microbial Observatories program (MCB-0702395), the Long Term Ecological Research program (NTL-LTER DEB-1440297), and an INSPIRE award (DEB-1344254), from NASA through project #NNA13AA94A administered by the NASA Astrobiology Institute, from the University of Wisconsin-Madison through Microbiome Initiative, and from the Deutsche Forschungsgemeinschaft (DFG, LA 4177/1-1). This material is also based upon work that supported by the National Institute of Food and Agriculture, U.S. Department of Agriculture (Hatch Project 1002996). The work conducted by the U.S. Department of Energy Joint Genome Institute, a DOE Office of Science User Facility, is supported by the Office of Science of the U.S. Department of Energy under Contract No. DE-AC02-05CH11231.

## FUNDING INFORMATION

U.S. NSF Microbial Observatories program (MCB-0702395) Katherine D McMahon

U.S. NSF Long Term Ecological Research program (NTL-LTER DEB-1440297) Katherine D McMahon

U.S. NSF INSPIRE award (DEB-1344254) Katherine D McMahon

U.S. National Institute of Food and Agriculture, U.S. Department of Agriculture (Hatch Project 1002996)

Katherine D McMahon

NASA Astrobiology Institute (NNA13AA94A) Eric E Roden

University of Wisconsin-Madison (Microbiome Initiative) Eric E Roden

Deutsche Forschungsgemeinschaft (DFG), Grant Number LA4177/1-1 Maximilian P Lau

## List of Supplemental Information

Supplemental Methods

Table S1. Lake characteristics

Table S2. List of MHCs that might be involved in EET in the metagenomes

Table S3. List of Cyc2-like genes in the metagenomes

